# HIGH THROUGHPUT QUANTITATION OF HUMAN NEUTROPHIL RECRUITMENT AND FUNCTIONAL RESPONSES IN AN AIR-BLOOD BARRIER ARRAY

**DOI:** 10.1101/2024.05.10.593624

**Authors:** Hannah Viola, Liang-Hsin Chen, Seongbin Jo, Kendra Washington, Cauviya Selva, Andrea Li, Daniel Feng, Vincent Giacalone, Susan T. Stephenson, Kirsten Cottrill, Ahmad Mohammed, Evelyn Williams, Xianggui Qu, Wilbur Lam, Nga Lee Ng, Anne Fitzpatrick, Jocelyn Grunwell, Rabindra Tirouvanziam, Shuichi Takayama

## Abstract

Dysregulated neutrophil recruitment drives many pulmonary diseases, but most preclinical screening methods are unsuited to evaluate pulmonary neutrophilia, limiting progress towards therapeutics. Namely, high throughput therapeutic screening systems typically exclude critical neutrophilic pathophysiology, including blood-to-lung recruitment, dysfunctional activation, and resulting impacts on the air-blood barrier. To meet the conflicting demands of physiological complexity and high throughput, we developed an assay of 96-well Leukocyte recruitment in an Air-Blood Barrier Array (L-ABBA-96) that enables *in vivo*-like neutrophil recruitment compatible with downstream phenotyping by automated flow cytometry. We modeled acute respiratory distress syndrome (ARDS) with neutrophil recruitment to 20 ng/mL epithelial-side interleukin 8 (IL-8) and found a dose dependent reduction in recruitment with physiologic doses of baricitinib, a JAK1/2 inhibitor recently FDA-approved for severe COVID-19 ARDS. Additionally, neutrophil recruitment to patient-derived cystic fibrosis sputum supernatant induced disease-mimetic recruitment and activation of healthy donor neutrophils and upregulated endothelial e-selectin. Compared to 24-well assays, the L-ABBA-96 reduces required patient sample volumes by 25 times per well and quadruples throughput per plate. Compared to microfluidic assays, the L-ABBA-96 recruits two orders of magnitude more neutrophils per well, enabling downstream flow cytometry and other standard biochemical assays. This novel pairing of high-throughput *in vitro* modeling of organ-level lung function with parallel high-throughput leukocyte phenotyping substantially advances opportunities for pathophysiological studies, personalized medicine, and drug testing applications.

## Introduction

Modulating neutrophilic inflammation has historically been challenging in respiratory medicine^1,2^ despite an expanding pool of immunomodulators available for clinical use^3–6^. A major bottleneck has been the lack of preclinical screening tools that adequately capture human-specific neutrophil responses such as priming while in the circulation, pathological recruitment into tissues, and the secondary acquisition of dysfunctional states therein^2,7–9^. Although animal models can capture complex tissue-level pathophysiology such as immune cell recruitment and activation^10,11^, their use is hindered by low throughput and most importantly, evolutionary divergence between species. Rodents, for example, lack interleukin-8 (IL-8), a central neutrophil chemoattractant in human disease^10,12,13^. As an alternative, *in vitro* platforms have been developed to support human neutrophil-targeted drug screening^14–18^. So far, however, technical demands and practical constraints of *in vitro* systems for human neutrophil-focused drug screening have limited their impact. Since candidate therapies aim to influence functional behavior of neutrophils, such as their recruitment, priming, activation, and signaling to resident tissue cells, these must be featured in screening models^2^. Consequently, transmigration systems are needed that incorporate disease-relevant stimuli and allow neutrophil engagement with both endothelium and epithelium to induce their physiological cascade of activation^19,20^. Such advanced transmigration models must be coupled to a practical downstream workflow that reduces cost and uses accessible technologies to deliver relevant, functional outcomes with high throughput^21,22^.

Microphysiological systems (MPS) use human cells in co-culture to capture emergent functional behaviors of multicellular systems, like immune cell recruitment, paracrine signaling loops, and mucosal barrier function^23,24^. MPS must balance throughput, robustness, physiologic relevance, efficient use of limited human cells and clinical samples, and the provision of quantitative assay outcomes^21,24^. Microfluidic “lung-on-a-chip” devices are prominent lung MPS that often feature bilayer epithelial-endothelial co-culture to mimic the alveocapillary (i.e., air-blood) barrier. Microfluidics can also apply physiologic forces like shear, strain, and compression to the air-blood barrier to modulate its properties^24–27^. However, trans-endothelial/epithelial immune cell recruitment assays in microfluidics are constrained. Microfluidics’ small size precludes recruitment of more than 10’s to 100’s of neutrophils per chip^16,28–30^, an insufficient amount for downstream phenotyping with reliable, established methods like flow cytometry. Performing non-terminal barrier measurements in microfluidics, such as trans-epithelial electrical resistance (TEER), requires custom device engineering^31^. Mechanical stress imparted microfluidic channels during neutrophil retrieval could affect their phenotype, skewing outcomes^32,33^. Finally, generating and culturing microfluidic lungs-on-a-chip typically requires bespoke equipment and advanced technical skills that are inaccessible outside of specialized laboratories, limiting their overall reach^34^. Under these limitations, the few microfluidic transmigration models reported have only measured the number of migrated cells and/or their rolling velocity, both using representative images from low-throughput microscopy^16,28–30,35,36^.

The Boyden chamber is a second well-known platform consisting of a porous plastic membrane that separates a culture well so that cells migrate from the top to bottom chamber. Typical assays, like Transwell^®^ and Alvetex^®^ filters in 12- to 24-well plate format, can recruit adequate neutrophils from the top to bottom chamber for downstream immunophenotyping^37–39^. However, 12- and 24-well assays require high neutrophil numbers (millions/well) and clinical specimen volumes (millimeters/well) that constrain the amount of conditions that can be tested with these limited, valuable materials. Additionally, the Alvetex^®^ recruitment assay requires manual inversion of the filter insert to place epithelial cells on the filter underside, a technically challenging, low-throughput procedure; additionally, Alvetex^®^ cannot accommodate bilayer endothelial culture^40^. A 96-well Transwell^®^ array compatible with high throughput screening (HTS) is commercially available (Corning, HTS Transwell^®^ 96-well Permeable Support), but bilayer co-culture on the Transwell-96 is hardly reported because underside cell seeding is prohibitively difficult at this miniscule filter size^41^. Cell-free and trans-endothelial recruitment in the Transwell-96 are reported more often^42,43^, but sacrificing either cell type is highly undesriable; both are critical participants in pulmonary neutrophil recruitment^44,45^. Taken together, the available neutrophil recruitment assays must compromise between clinical sample efficiency, throughput, physiologic relevance, ease of use, and availability of downstream analyses. These limiting compromises motivate novel approaches to meeting these conflicting technical demands to advance studies of neutrophilic inflammation.

Therefore, we report a novel assay, termed the 96-well Leukocyte recruitment in an Air-Blood Barrier Array (L-ABBA-96), that precisely quantifies neutrophil number and activation status before and after endothelial-to-epithelial recruitment in a miniature, high throughput format coupled to automated flow cytometry for immunophenotyping. The L-ABBA-96 builds off of our recently described method for underside epithelial seeding and bilayer co-culture with endothelial cells on the Transwell-96^41^. Healthy donor neutrophils placed in the endothelial chamber extravasate physiologically through the air-blood barrier toward chemoattractants in the epithelial chamber. We show that neutrophils recruited to IL-8 become activated by exposure to the epithelial milieu and acquire a phenotype similar to airway neutrophils *in vivo*. Interestingly, non-recruited endothelial neutrophils also acquire a physiologically relevant phenotype, mimicking the priming seen in circulating neutrophils. We then show that the immunomodulator baricitinib, JAK1/2 inhibitor recently FDA-approved to treat COVID-19-associated inflammation^46^, reduces neutrophil recruitment and shifts activation marker expression for both air- and blood-side neutrophils^47^. Finally, the L-ABBA-96 recapitulates cystic fibrosis inflammation; epithelial-side patient-derived sputum caused upregulation of e-selectin shifted surface markers to CF-mimetic phenotypes in both recruited and non-recruited neutrophils.

In summary, the L-ABBA-96 substantially improves the physiologic relevance of convenient and accessible plate-based transmigration assays and increases the volume of information gained from them by enabling traditional immunophenotyping and precise, high-throughput quantitation with automated flow cytometry. Accessibility of highly-physiological and high-throughput assays to those not specialized in bioengineering is critical to advancing the field’s understanding of pulmonary inflammation and development of neutrophil immunomodulators.

## Results

### IL-8 and LTB4 dose-dependently recruit primary human neutrophils in L-ABBA-96

Neutrophil recruitment from the circulation is a key event that sets the stage for tissue responses early after injury or infection. Chemotaxis and transmigration, as well as soluble signals, prime quiescent circulating neutrophils for their eventual antimicrobial and pro-tissue regenerative functions once they arrive at the injury site^19,48,49^. Priming is followed by activation during transmigration, *via* engagement with endothelial, interstitial, and epithelial ligands; sensing of chemoattractants, extracellular matrix and cytokines; and experiencing mechanical forces while squeezing through tissue^32,50,51^. Once reaching the site of injury, activated neutrophils release antimicrobial effectors, growth factors, cytokines, chemokines and exosomes that influence the tissue microenvironment^52^,. Therefore, rather than studying neutrophils isolated from blood, incorporating the key activating process of transmigration is critical to understanding neutrophilic inflammation and designing effective, targeted therapeutics to modulate it. The L-ABBA-96 aims to recapitulate these key *in vivo* steps of neutrophil priming, recruitment, and activation in a high-throughput *in vitro* platform (**Figure 1A, B**). Airway epithelial cells are seeded on the underside of Transwell membranes as previously described^41^, opposite a confluent human endothelium. Chemoattractant is added to the lower epithelial compartment, and neutrophils are added to the upper endothelial compartment. Over several hours, neutrophils transmigrate through the endothelium, the collagen-coated Transwell membrane, and the air-liquid interface-differentiated epithelium to reach the chemoattractant.

**Figure 1.**
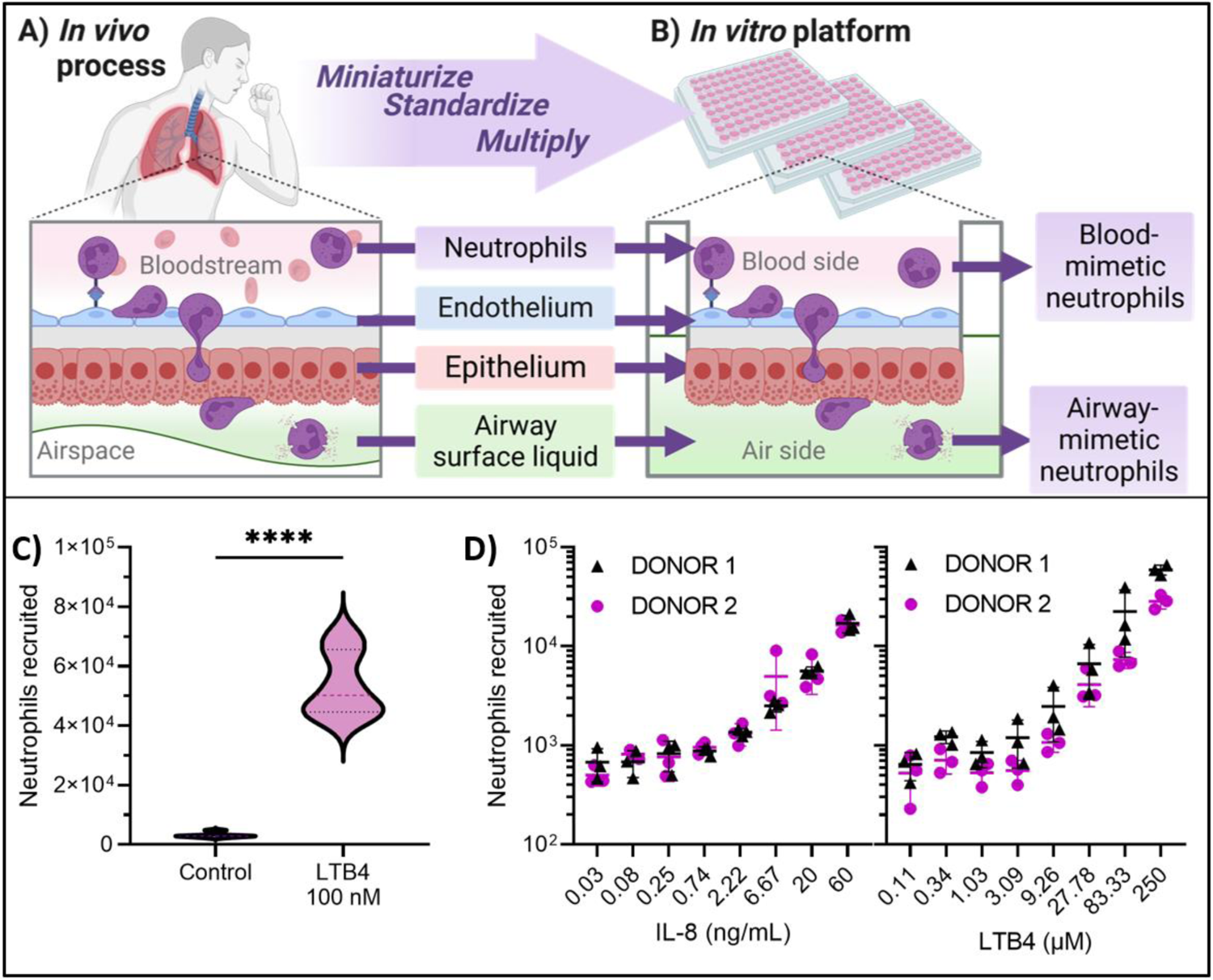
A miniaturized air-blood barrier array recapitulates transendothelial-transepithelial neutrophil recruitment in vitro. Neutrophil recruitment to the distal lungs through the endothelium and then epithelium (A) is recapitulated in a miniaturized, standardized platform in 96-well format (B) using off-the-shelf reagents and standard cell culture equipment. C) Despite miniaturization, neutrophils are recruited in high numbers with a high degree of specificity between negative and positive control conditions. n=6 per condition (3 replicates per donor, 2 donors). Compared with student’s t-test.****p<0.0001. D) Neutrophils are dose-dependently recruited to IL-8 and LTB4, two central chemoattractants in human lung diseases. The assay’s high sensitivity and many replicates enable detection of a difference in the response to LTB4, but not IL-8, between two donors, according to two-way ANOVA (p<0.0001). n=6 per condition (3 replicates per donor, 2 donors). Figure created with BioRender software.

To establish quality control for transmigration assays, we optimized and standardized the L-ABBA-96 culture protocol to minimize edge position effects, a frequent concern with plate-based assays^53^. Controlling temperature and humidity prevented edge position effects on barrier strength, as measured by trans-epithelial/endothelial electrical resistance (TEER) (**Supplemental Figure 1, Supplemental Method 1**). Plates with an average TEER ≥200 Ω*cm^2^ showed a negligible correlation between air-blood barrier strength in the numbers of recruited neutrophils (**Supplemental Method 2**; **Supplemental Figure 2-3**). With standard operating procedures and quality control metrics in place, we first analyzed recruitment of primary, human circulating neutrophils to increasing doses of the chemoattractants IL-8 and LTB4 (Figure 1D). The assay’s throughput accommodates an 8-fold dose-response curve with multiple donors, replicates, and chemoattractants, which minimizes experimental batch effects and improves the assay’s resolution. We detected a significant difference between two independent donors in the dose-response curve for LTB4 (p<0.0001), but not IL-8 (p=93.97) using a two-way ANOVA for the effects of chemoattractant and donor on the number of recruited neutrophils. While the effect is minor and secondary to the demonstration of precise and controlled dose-responses in the assay, it is interesting to note that the human LTB4 receptor is exceptionally epigenetically variable^54^ and could cause biological variability in neutrophil responses between donors that are captured by the L-ABBA-96.

### L-ABBA-96 mimics activation marker shifts induced by lung neutrophil recruitment

The L-ABBA-96 platform recovers sufficient neutrophils for flow cytometric analysis (Gating scheme is shown in **Supplemental Figure 4**) for both unmigrated (top chamber) and migrated (bottom chamber) cells (Figure 2A). Thus, it enables evaluation of surface marker expression at 3 critical stages: (i) on freshly blood-isolated neutrophils immediately before placement in the assay; (ii) on unmigrated neutrophils (endothelial side); and (iii) on migrated neutrophils (epithelial side) accumulated at the air-blood barrier for the duration of the assay. We compared surface expression of key neutrophil phenotypic markers CD62L, CD16, CD66b, and CD63 after 14-hour exposure of endothelial-side neutrophils to epithelial-side IL-8 (20 ng/mL), between 3 independent donors. Similar neutrophil numbers were recruited across the 3 donors (p=0.19, one-way ANOVA with post-hoc Tukey’s t-test) (Figure 2B)). For all donors, neutrophil surface expression of CD62L, CD16, CD66b, and CD63 were consistent with quiescent, primed and activated phenotypes of blood, unmigrated, and migrated neutrophils, respectively. For each marker, differences were assessed by two-way ANOVA between donor and condition with post-hoc Tukey’s t-test for main row effect (i.e., averaged donors). CD62L is shed by enzymes during neutrophil priming^48^ and further by ectodomain shedding by endothelial and epithelial disintegrins during transmigration^55,56^ in line with our observations (Figure 2Di). CD16 (FcγRIIIb) can be shed during priming, activation, and chemotaxis^57–59^, but it is also mobilized from intracellular stores during transmigration^59,60^. Our assay suggests unmigrated neutrophils were primed, reflected by decreased CD16, and then recovered expression with the stimulus of transmigration (Figure 2Dii). CD66b and CD63, indicative of secondary and azurophilic degranulation, respectively, were only increased in the migrated neutrophils, suggesting migration stimulates activation and subsequent degranulation (Figure 2Diii**-iv**)^8^.

**Figure 2.**
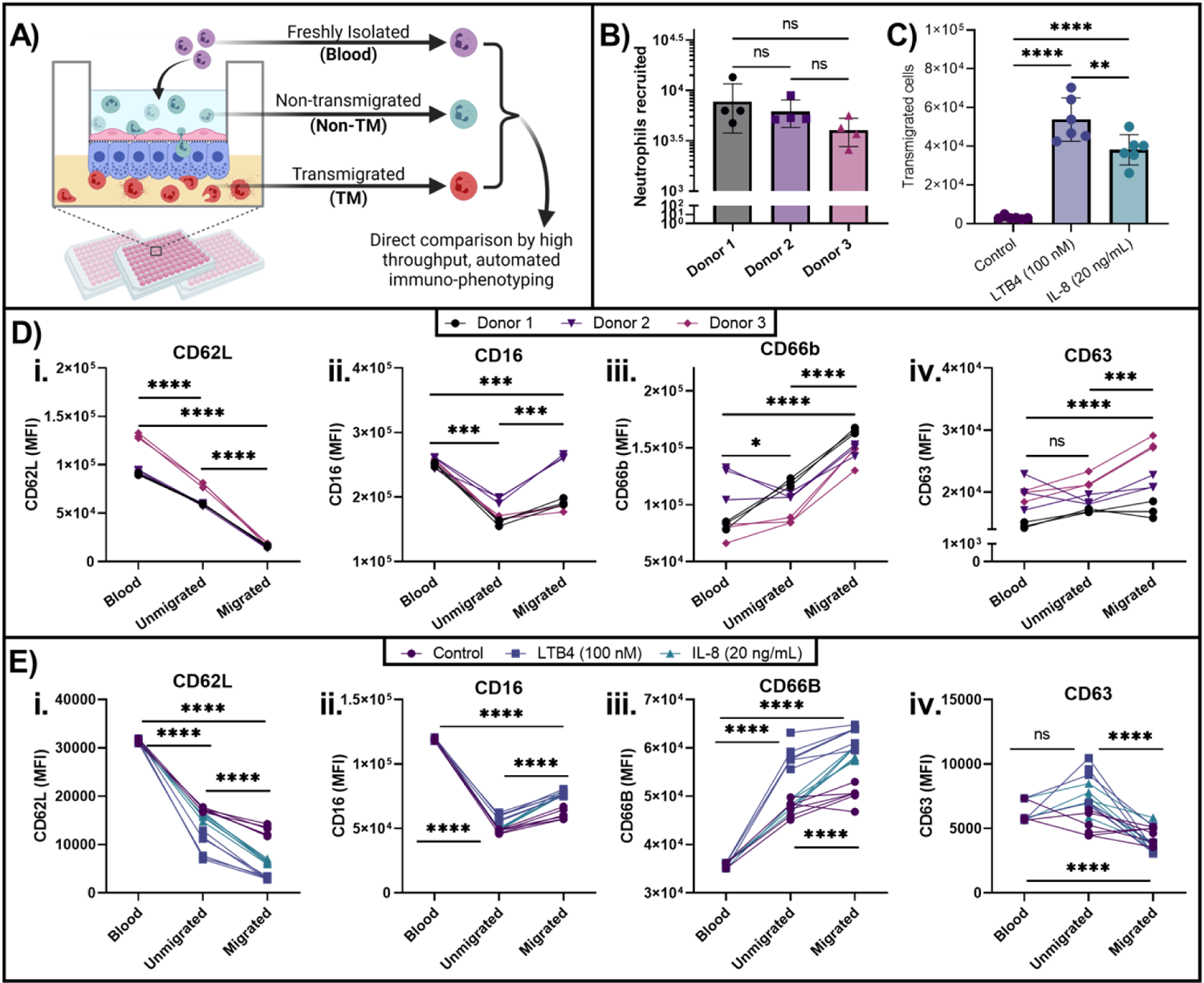
Neutrophil recruitment in vitro reflects physiologic shifts in key surface markers. **A)** Key activation markers CD62L, CD66b, CD16, and CD63 were compared between blood, unmigrated, and migrated neutrophils across 3 donors for 1 chemoattractant **(B, D)** and 3 chemoattractant conditions for 1 donor **(C, E)**. **B, D)** Neutrophils were recruited to epithelial IL-8 (20 ng/mL), LTB4 (100 nM), or no chemoattractant for 14 hours (n=4-6 replicates/donor). **B)** Comparable numbers of neutrophils from each donor are recruited to IL-8 (20 ng/mL). **C)** LTB4 (100 nM) attracts more neutrophils than IL-8 (20 ng/mL). **Di, Ei)** CD62L, or L-selectin, is shed during priming, and further by transmigration, due to engagement with endothelial and epithelial disintegrins, across donors and recruitment conditions. **Dii, Eii)** CD16 (FcγRIII) is shed during priming and activation of neutrophils, as in the non-TM population, but can be replaced by intracellular reserves after transmigration, which is consistent with the increase seen in TM neutrophils in our assay for multiple donors and chemoattractants. **Diii, Eiii)** CD66b is a human-specific marker of neutrophil degranulation associated with activation. Physiologic recruitment to the lung increases CD66b and this is captured in our assay across donors and chemoattractants. LTB4 activates non-recruited neutrophils more than IL-8. **Div, Eiv)** Degranulation of azurophilic granules, represented by CD63, indicates neutrophil activation, which is mildly induced in certain donors by recruitment to large concentratrations of chemoattractant, such as the 20 ng/mL IL-8 used here. Interestingly, CD63 was not increased for the single donor across conditions, suggesting donor effects. B, D) N=3 donors, n=4 replicates/donor; **C, E)** N=1 donor, n=3 replicates/donor. **D, E)** Analyzed by two-way ANOVA with post-hoc Tukey’s t-test (main row effect). *p<0.05, **p<0.01, ***p<0.001, ****p<0.0001. A) was created with BioRender.

**Figure 3.**
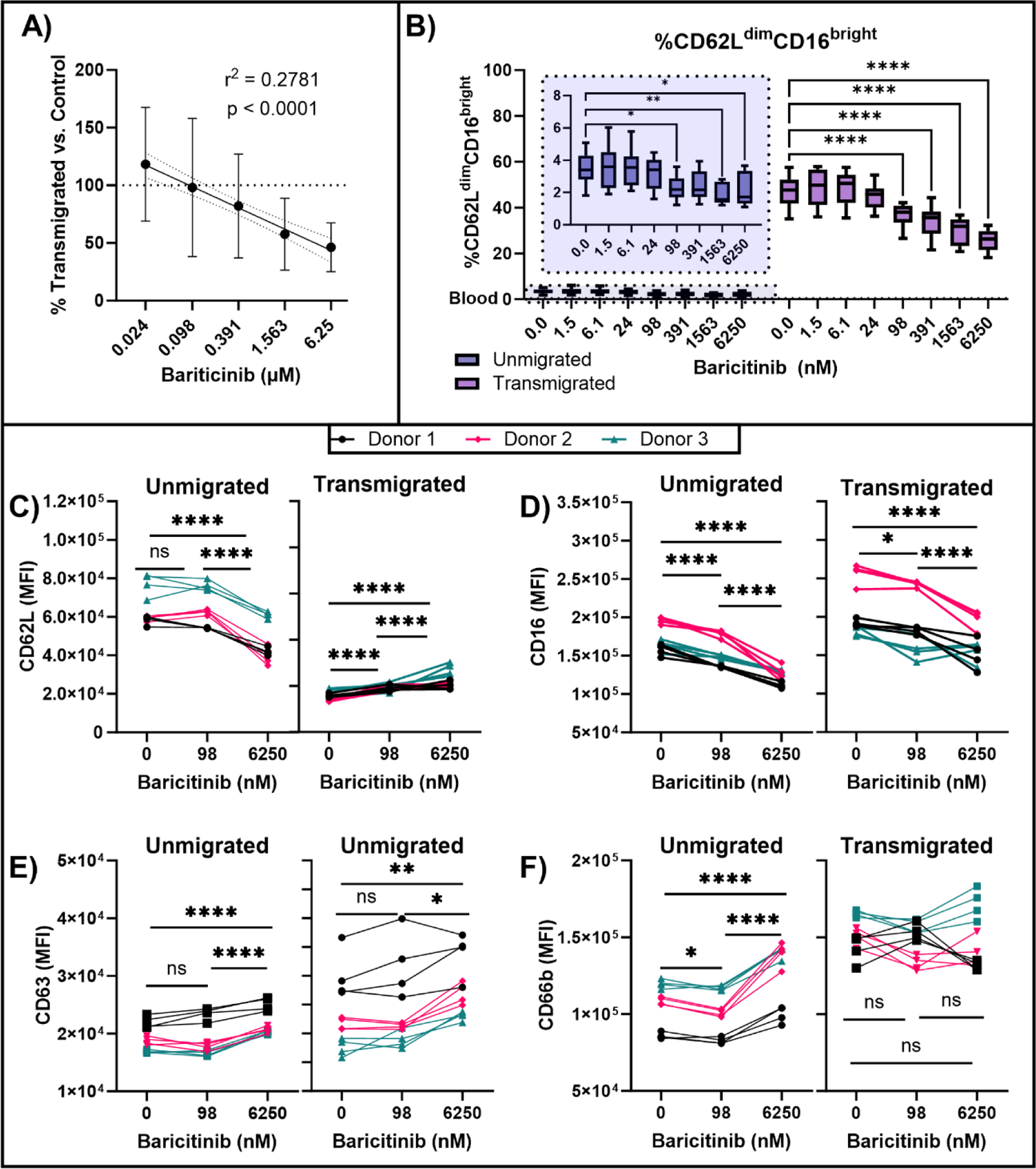
Baricitinib on the endothelial side of the L-ABBA-96 dose-dependently modulates neutrophil phenotype. **A)** Across 5 donors and 9 assays, baricitinib significantly reduced neutrophil recruitment towards IL-8 (20 ng/mL). A simple linear regression of the linearized data revealed a weak (r^2^=0.2781) yet highly significant (p<0.0001) negative correlation between drug concentration and recruited neutrophil number. N=5 donors, n=3-6 replicates/donor/assay. **B)** Highly activated CD62L^dim^CD16^bright^ neutrophils are attenuated by baricitinib on the blood side (inset, repeated measures one-way ANOVA with Dunnett’s post-hoc test) and the air side (two-way ANOVA with post-hoc Dunnett’s test against zero drug control, matched by well for TM and non-TM). N=3 donors, n=4 replicates/donor. **C-F)** A non-physiologically relevant dose of 6250 nM baricitinib induced mild priming and activation that was negligible at a clinically relevant dose of baricitinib (98 nM). Two-way ANOVA with post-hoc Tukey’s t-test (main row effect) Within unmigrated or transmigrated,. *p<0.05, **p<0.01, ***p<0.001, ****p<0.0001.

These observations suggest that the L-ABBA-96 captures key features of the physiologic cascade of neutrophil recruitment in the lung^61^, wherein neutrophils are quiescent in the bloodstream, primed during endothelial arrest and/or exposure to certain inflammatory signals^62^, and activated after transmigration to the lumen^61^. Interestingly, while neutrophil recruitment to IL-8 did not reveal a statistically different donor effect on neutrophil number (similar to in Fig 1), flow cytometric quantitation of activation markers appear to reveal activation phenotype-based donor effects. According to two-way ANOVA analyzing the effect of transmigration (blood-isolated, unmigrated, or migrated) and donor-dependent biological variability in neutrophils, donor variation accounted for 23.6 ± 19.3% of variation, on average, in the expression of surface markers (**Supplemental Tables 1-2**). Transmigration condition (blood-isolated, unmigrated, migrated) accounted for the majority (62.1 ± 25.3%) with minor interaction effects comprising the remainder (10.7 ± 4.4). These results underscore our platform’s ability to monitor neutrophil through various steps of priming and activation that accompany the transmigration process.

We also analyzed response of neutrophils from a single donor to IL-8 (20 ng/mL), LTB4 (100 µM), or control (epithelium submerged in media with no additional chemoattractant) using the same activation markers (Figure 2E). Transmigration effects were statistically significant (p<0001, two-way ANOVA) for all markers except CD63 (p=0.1548), which was not increased by recruitment to any chemoattractant for this particular donor. Taken together, the L-ABBA-96 can consistently induce neutrophil priming, recruitment, and activation and quantify the changes with sufficient resolution, consistency, and number of replicates that biological variability of response from different donor neutrophils or chemoattractants can be discerned.

### Baricitinib dose-dependently inhibits neutrophils’ recruitment and modulates their surface phenotype

We then evaluated the influence of baricitinib, a small-molecule JAK1/2 inhibitor, on IL-8-driven neutrophil recruitment in a model of severe pulmonary inflammation^63^. Baricitinib was recently approved by the U.S. Food and Drug Administration for controlling pulmonary inflammation in severe Coronavirus Disease 2019 (COVID-19) and is a promising immunomodulator for many other inflammatory lung diseases^46,63–67^. We used IL-8 alone at 20 ng/mL, a high concentration similar to that in bronchoalveolar lavage fluid from patients with severe COVID-19^68,69^. Figure 3A shows log-log plots of number of neutrophils vs. the baricitinib dose for a combined N=5 donors across 9 assays, demonstrating a small (r^2^=0.2781) but robust (p<0.0001) linearized dose-dependent reduction in recruited neutrophils (Figure 3A). Interestingly, reduced neutrophil recruitment was also observed in rhesus macaques treated with baricitinib for COVID-19^70^ but was not observed in a recent *in vitro* IL-8 transendothelial migration study using larger Transwell inserts^71^, supporting our assay’s unique sensitivity and physiologic relevance. Additionally, the half maximal inhibitory concentration (IC50) of baricitinib varied widely across 3 neutrophil donors tested on the same L-ABBA-96 assay plate (1.409 µM, 1.86 µM, and 95.9 nM) (**Supplemental Figure 5**). These IC50 values may be explained by the fact that IL-8 mediates chemotaxis through JAK3 rather than baricitinib’s main targets JAK1/2^72^. Cell-free assays determined baricitinib’s IC50 against JAK1 and JAK2 as 5.9 nM and 5.7 nM, respectively, and ∼420 nM for JAK3^63^, which is within one order of magnitude of our measured IC50s. Interestingly, maximum plasma concentration of baricitinib is 97.5 ± 21.6 nM (36.2 ± 8.0 ng/mL)^73^, close to the IC50 determined for one donor in the L-ABBA-96 (95.9 nM).

Additionally, a population of highly active, immunosuppressive neutrophils with a CD62L^dim^CD16^bright^ phenotype has recently been identified in COVID-19 and other inflammatory conditions^74–77^. In the L-ABBA-96, baricitinib dose-dependently reduced the proportion of IL-8-induced CD62L^dim^CD16^bright^ neutrophils in both unmigrated and migrated populations (**Supplemental Figure 6**). Conversely, with increasing baricitinib dose, CD62L was shed more in unmigrated neutrophils and CD16 was shed more in both unmigrated and migrated neutrophils. This suggests that baricitinib may enhance priming of unmigrated neutrophils, through an as yet unknown mechanism. Interestingly, CD66b and CD63 were not significantly impacted in either unmigrated or migrated neutrophils, suggesting no significant degranulation is induced until a dose of baricitinib of 6.25 µM, which is 2 orders of magnitude greater than its reported maximal serum concentration of around 100 nM^73^. The clinically relevant 98 nM dose in our study was enough to significantly reduce the proportion of CD62L^dim^CD16^bright^ neutrophils among unmigrated neutrophils without inducing appreciable activation (Figure 3C-F). Taken together, baricitinib in the L-ABBA-96 dose-dependently attenuates neutrophil recruitment and induction of the pathological CD62L^dim^CD16^bright^ phenotype.

### Recruitment to cystic fibrosis sputum-derived airway supernatant induces disease-mimetic inflammatory phenotypes

To demonstrate the capability of the L-ABBA-96 to capture disease-specific pathophysiology, the L-ABBA-96 epithelium was exposed to pooled human cystic fibrosis (CF) sputum-derived airway supernatant^37^ (CFASN) (Figure 4A). Previous Alvetex-based recruitment experiments by our group show that trans-interstitial and epithelial migration into CFASN programs neutrophils towards disease-characteristic features of granule-releasing, immunomodulatory and metabolically active (“GRIM”) cells, with low bacterial killing^37^. GRIM neutrophils present in the CF lung lumen are CD62L^dim^ due to transmigration^56^, CD66b^bright^ and CD63^bright^ due to high release of secondary and primary granules (the latter containing neutrophil elastase), and simultaneously CD16^dim^ due cleavage by elastase^37^. Here, in the L-ABBA-96, neutrophils were allowed to transmigrate for 18 hours into CFASN (or IL-8 at 20 ng/mL as a control), diluted in cell culture medium at a ratio between 1:12 to 1:1, as specified. Neutrophils transmigrated dose-dependently to increasing concentrations of CFASN. Interestingly, TEER increased and decreased at low and high CFASN concentrations respectively (Figure 4A). The endothelium also upregulated E-selectin following exposure to CF-ASN in the epithelial compartment in a dose-dependent manner (Figure 4B). Consistently, CF patient serum contains elevated soluble E-selectin^78,79^.

**Figure 4.**
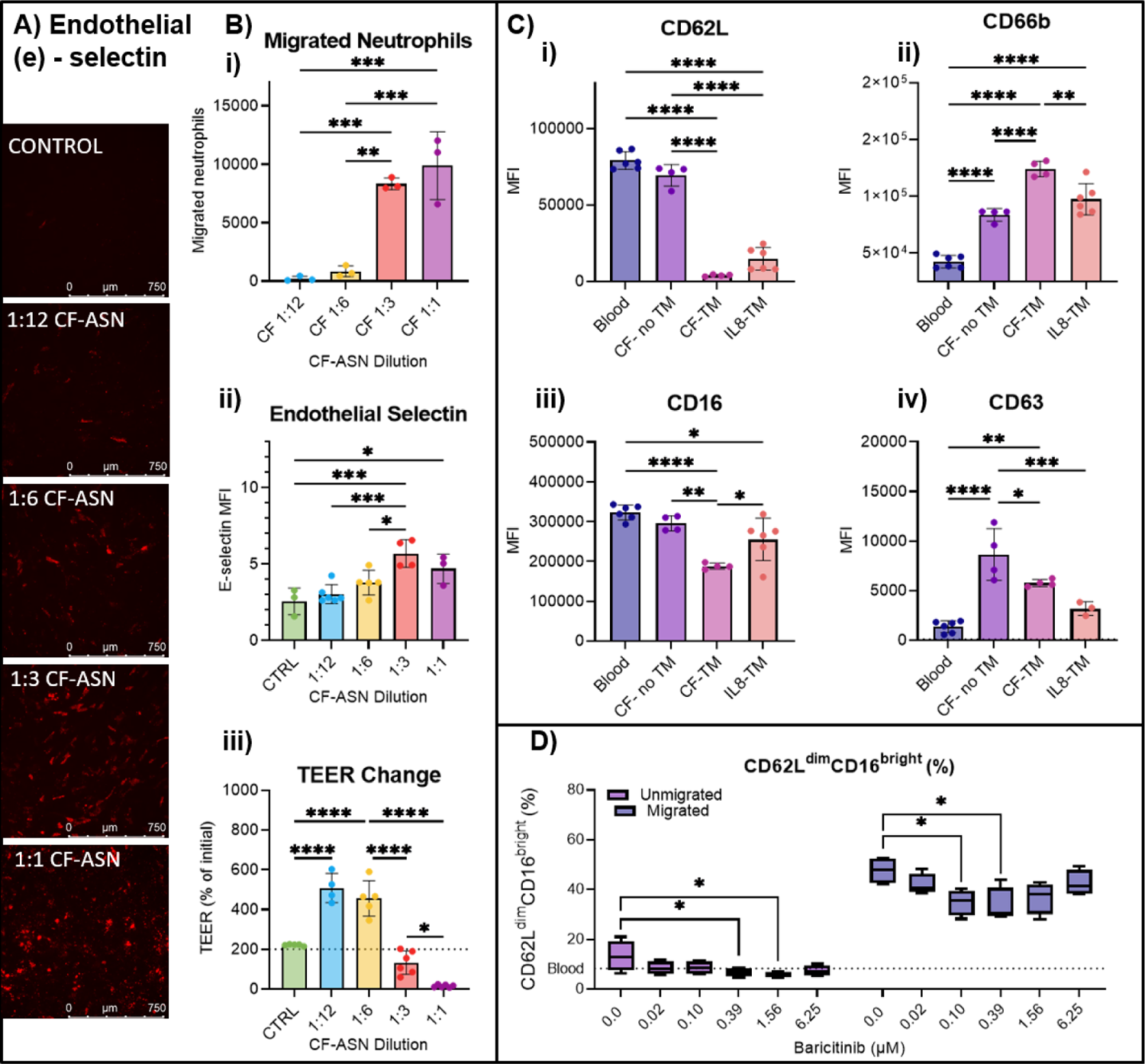
L-ABBA-96 recapitulates pulmonary neutrophilia and elevated blood neutrophil activation characteristic of cystic fibrosis. Healthy donor neutrophils were recruited towards pooled, patient-derived CF-ASN containing the soluble factors from sputum without mucus or cells. Lung-recruited CF neutrophils demonstrate significant activation, and characteristically lose CD62L and CD16 while gaining CD66b and CD63. A) Endothelial selectin (e-selectin) was upregulated on endothelial cells in the L-ABBA-96 after 18 hours’ exposure to epithelial-side CF-ASN (quantified in Bii)). Bi) Neutrophils migrate dose-dependently to CF-ASN Bii) E-selectin protein is upregulated dose-dependently by CF-ASN on the epithelium according to quantitative immunofluorescence. Biii) Epithelial-endothelial barrier strength, measured by TEER, is compromised by high concentrations of CF-ASN. C) Neutrophils recruited to CF-ASN (hereafter CF-TM neutrophils) display more activation than those recruited to IL-8 (IL8-TM). Ci, Cii, Ciii, Civ) CF-TM neutrophils trend toward losing more CD62L and gaining more CD63 than IL8-TM. They statistically lose more CD16 and gain more CD66b than IL8-TM. D) Unmigrated and migrated neutrophil proportion of CD62L^dim^CD16^bright^ is dose-dependently reduced by baricitinib. All except D) were analyzed by one-way ANOVA with post-hoc Tukey’s t-test. D) Two-way ANOVA with post-hoc Tukey’s t-test. C) 2 donors, 2-3 replicates/donor; D) 2 donors, 2 replicates/donor. *p<0.05, **p<0.01, ***p<0.001, ****p<0.0001.

Next, we evaluated the mobilization of surface markers (Figure 4C) on neutrophils from the CFASN-treated L-ABBA-96. We excluded the 1:12 and 1:6 conditions due to low transmigrated cell numbers. At the higher concentration CFASN conditions, the transmigrated neutrophils shed CD16 and gained CD66b significantly more than neutrophils recruited to IL-8 in the same assay. Recruited CF neutrophils also trended towards lower CD62L and higher CD63 than recruited IL-8 neutrophils, although significance was not achieved by one-way ANOVA^80^. Overall, the greater activation indicated by the surface markers measured is consistent with a CF-like activation of recruited neutrophils.

Interestingly, unmigrated (blood-side) neutrophils also displayed a CF-like phenotype in the L-ABBA-96. CF patients have an elevated proportion of circulating CD62L^dim^CD16^bright^ neutrophils^81^, which was reflected in the CF-ASN exposed L-ABBA-96 (Figure 4D). As previously mentioned, this population has been characterized as immunomodulatory and highly activated in acute inflammation, but their role in CF is not well characterized^74,82^. CF patient circulating neutrophils are approximately 20% CD62L^dim^CD16^bright^; in the L-ABBA-96, a similar proportion (13.2 ± 5.3%) of neutrophils were CD62L^dim^CD16^bright^ despite originating from healthy donors and failing to develop this magnitude of a response to IL-8 (20 ng/mL) during the same assay^81^. Further, this population on both air and blood sides were reduced by clinically relevant doses of baricitinib, mirroring results from Figure 3 for recruitment to IL-8. In summary, the L-ABBA-96 recapitulates multimodal features of CF pulmonary inflammation, including neutrophilia, e-selectin upregulation, significant recruitment-induced activation, and the appearance of a blood-side population of CD62L^dim^CD16^bright^ neutrophils at a similar proportion to human patients, supporting its value for disease-specific investigations.

## Discussion

Current methods for *in vitro* assessment of human neutrophil-driven lung inflammation are limited in throughput, standardization, phenotypic relevance, or ability to recover sufficient numbers of cells to perform high-content phenotypic analyses. Here, we bridge this gap with a high-throughput, miniaturized assay comprised of an array of leukocyte-laden air-blood barriers, the L-ABBA-96, that can be coupled with multifactorial outcome quantification of airway barrier function and neutrophil phenotype by flow cytometry. Quantification of these parameters across more than a thousand air-blood barriers demonstrate reproducibility and consistency across different wells, plates, neutrophil donors as well as versatility across a variety of stimulants including chemoattractants, cytokines, and patient fluid samples.

The L-ABBA-96 benefits from operating at the meso-scale, between microfluidics and traditional plate-based assays. This allows recovery of adequate numbers of cells (>1,000’s transmigrated cells) from each well of 96-well format plates in a manner compatible with high-throughput flow cytometry while being volume efficient (< 100 µL per bottom well) to minimize use of clinical specimens and costly reagents. Indeed, most plate-based assays do not offer tissue-level complexity and miniaturization, as these features are typically reserved for microfluidic devices. As opposed to microfluidics, however, our platform requires similar technical complexity, skill, and equipment to existing plate-based transmigration assays that are widely used. Our platform also excludes the use of glucocorticoids commonly used to stimulate epithelial differentiation^83^. Therefore, we are able to study inflammation-modulating therapies without interference.

Detailed dose-response studies of neutrophil recruitment and activation across multiple donors is uncommon, at least in part, due to technical limitations such as low throughput or limited numbers of transmigrated cells hampering downstream analyses. The L-ABBA-96 accommodates 8-fold dose-response curves on a single plate with many replicates to increase confidence while reducing experimental variability. Each replicate has 3 interacting cell types, epithelial-side exposure to chemoattractants or clinical samples, the endothelial-side is dosed with different concentrations of a therapeutic, and outcomes quantified through TEER and flow cytometry of unmigrated (blood-side) and transmigrated (airway-side) neutrophils. The procedures are amenable to scale-up and automation using off-the-shelf cultureware, liquid handlers, and measurement systems including underside epithelial seeding, leukocyte loading and transmigration, barrier property measurement, and flow cytometry.These unique advantages position the L-ABBA-96 to characterize pulmonary insults such as inhalants from cigarettes, e-cigarettes and other electronic nicotine delivery systems using novel immune-relevant outcome measures that may relate to pathophysiology.

Another key feature of the L-ABBA-96 is its ability to recover unmigrated neutrophils that remain on the endothelium but can nonetheless alter their phenotype in response to epithelial side stimulation. Interestingly, blood-side neutrophils in the L-ABBA-96 developed a CD62L^dim^CD16^bright^ phenotype reported as pathologic in systemic inflammatory conditions such as endotoxin challenge^74^ and cancer^88^. Most recently, an elevated CD62L^dim^CD16^bright^ neutrophil count in COVID patients was associated with developing pulmonary embolism^76^. Unexpectedly, in our assay baricitinib dose-dependently reduced the proportion of CD62L^dim^CD16^bright^ neutrophils among unmigrated and migrated neutrophils at clinically relevant doses. A recent case study reported the same effect of baricitinib on neutrophils in a COVID patient ^47,89^.

Compared to sophisticated lung-on-a-chip systems, the L-ABBA-96 has limitations in lacking physiological flow over the endothelium, pathophysiological fluid mechanical stress on the epithelium, as well as stretch. Compared to the Alvetex system that has a thicker interstitium with a complex geometry to represent later stage disease and promote neutrophil activation by collagen interactions and autocrine signaling, the L-ABBA-96 only has a thin collagen-coated porous membrane as the interstitium. Additionally, epithelial and endothelial cell types could be primary human lung-derived cells rather than cell lines or pooled umbilical vein cells. Beyond the lung, other mucosa-blood barriers that experience pathological neutrophil recruitment (e.g., the gut, skin, and oral, nasal mucosae) can be modeled in the L-ABBA-96 by exchanging the epithelial cells, expanding the platform’s impact.

Regardless, the L-ABBA-96 has strengths in using commercially available cultureware, readily available tools and reagents, and interfacing seamlessly with common automated flow cytometers making it widely accessible and immediately adaptable by all for further applications and improvements. While the biological findings of the work presented serve to demonstrate the capabilities and disease relevance of our platform, they also represent a significant technological advance for lung inflammation studies.

## Materials and Methods

### Preparation of Air-Blood Barrier Array (ABBA)

The ABBA was prepared in a standardized protocol as previously described by our group^41^. Briefly, we seeded human small airway-like epithelial cell line NCI-H441 (American Type Culture Collection (ATCC)) on the underside of the Transwell^®^ membrane (Corning HTS Transwell^®^ 96-well Permeable Support, polycarbonate, 3 µm pore size, #3386) using flotation. The endothelium was modeled with primary human umbilical vein endothelial cells (HUVECs, ATCC) that were pooled from multiple donors to reduce donor-specific variation.

### Neutrophil Transmigration Assays

Human peripheral blood was obtained through venipuncture under Georgia Tech Institutional Review Board (IRB)-approved protocols and isolated by negative selection according to manufacturer instructions (Miltenyi Biotec 130-104-434, 130-098-196). Trans-epithelial electrical resistance was measured as previously described with the EVOM2 and STX HTS for Corning 96 (World Precision Instruments)^41^. Chemoattractants (leukotriene B4, Cayman Chemical #20110; Interleukin 8, MyBioSource #MBS9718666) were prepared at the specified concentrations in ALI medium as described previously^41^. In CFASN assays, pooled patient airway supernatant obtained under an Emory IRB-approved collection was prepared as described^40^, diluted in ALI media to various concentrations as specified, and placed in the receiver plate. In drug testing assays, isolated neutrophils were suspended in media containing specified drug concentrations (baricitinib, Cayman Chemical #16707). Media was completely removed from the top chamber of the Transwell inserts and neutrophils suspended in ALI medium were placed in the top chamber of the Transwell at 100 uL/well, with 225,000-330,000 cells/well depending on the experiment. Neutrophils were placed in the top chamber and inserts were placed immediately into pre-warmed receiver plate containing the chemoattractant or patient specimen. The plate was incubated for 16 hours at 37 °C, 95% humidity, 5% CO2.

### Flow Cytometry

Freshly isolated neutrophils were stained for flow cytometry immediately after isolation in a FACS tube. Transmigrated neutrophils were collected into non-tissue culture treated round-bottom 96-well plates, blocked with Human TruStain FcX™ (Fc block, compatible with anti-CD16 staining) and stained according to optimized panels (**Supplemental Table 3**). Flow cytometry was performed using a Cytoflex S (Beckman Coulter) at 30 μL/min. Compensation was collected using single-antibody stained Abc Beads for antibodies and Arc beads for live/dead according to manufacturer instructions. Gating and compensation matrix was performed in FlowJo (BD Biosciences). Cell quantitation and median fluorescence intensities (MFI) for given markers were calculated using FlowJo. Gating strategy is described in **Supplemental Figure 3**.

### Statistical Analysis

All statistics were performed in GraphPad Prism 9. All significance tests were performed with α=0.05 (95% confidence). Figure 1C) Student’s t-test (unpaired). Figure 2B) Ordinary one-way ANOVA with post-hoc Tukey’s t-test. Figures 2C-2F) Two-way ANOVA with post-hoc Tukey’s t-test for main row effect (i.e., averaged donors). Figures 2F-2G) Two-way ANOVA with post-hoc Tukey’s t-test. Figure 3A**) (left)** One-way ANOVA with post-hoc Tukey’s t-test. Figure 3A**) (right)** Simple linear regression with 95% confidence intervals. Figure 3B**) (inset)** Ordinary one-way ANOVA with post-hoc Tukey’s t-test, Figures 3B**, 3Ci-iii, 3Di-iv)** Two-way ANOVA with post-hoc Dunnett’s t-test against control. Figures 4Cii**-iii, 4Di-iv)** Ordinary one-way ANOVA with post-hoc Tukey’s t-test. Figure 4Dv) Two-way ANOVA with post-hoc Dunnett’s t-test against control. For all graphs and charts, *\*p<0.05, **p<0.01, ***p<0.001, ****p<0.0001*.

## Acknowledgments

We thank blood donors who participated in this study, and the many other individuals who have contributed directly or indirectly to its implementation. This study was funded by NIH R01HL136141, U.S. Food and Drug Administration (FDA) Center for Tobacco Products (CTP) (75F40122C00146), EPA Grant #84045201, BME COVID Seed grant, and Georgia Tech EPICenter to ST; 5T32EB006343-08 to HV; NIH R01HL158058 and Cystic Fibrosis Foundation TIROUV22G0 to RT; K24 NR018866 to AF; and 5K23HL151897-04 to JG. We thank the GT IBB Core facilities for assistance with flow cytometry.

## Supporting Information for

### Supplemental Method 1: Humidity chamber

Evaporation gradients are considered a primary cause of edge effects because the relative humidity of most cell culture incubators averages at 95% and transiently drops further when the door is opened. The ABBA is particularly susceptible to evaporation due to the small media reservoir (<150 uL/well) and the air-exposed epithelium. Therefore, we developed a custom humidity chamber composed of a Nunc-well plate surrounded by 50-mL reagent reservoirs containing sterile water-saturated gauze with the plate in the center (supplementary Figure 1). ABBA plates that were cultured inside this humidity chamber within a standard cell culture incubator had reduced edge effects compared to plates without the humidity chamber (unpublished observation). Therefore, ABBA plates used in this study were cultured in the humidity chamber. Temperature gradients also cause edge effects including uneven cell seeding and growth rates due to the development of natural convection currents, which we observed when we performed seeding without temperature control. Therefore, all handling of the ABBA plate during seeding, culture, and assay preparation were performed on an electric cell culture heating apparatus with all reagents kept at 37 °C. Performing epithelial and endothelial seeding with pre-warmed plates on a heating plate significantly decreased barrier strength-related edge effects (unpublished observation).

### Supplemental Method 2: Edge Effects

Next, we investigated the influence of barrier strength and edge position on the number of migrated neutrophils, a key assay outcome. We hypothesized that wells with significantly low TEER compared to the other wells on the same plate would allow the passage of more neutrophils by allowing neutrophils to migrate through areas that were not completely covered with epithelial cells, consistent with the same observations made by Kidney and Proud (2000) during bronchial epithelial transmigration^1^. We conducted five independent transmigration experiments with assay plates having variable mean TEER (Table 1). Despite rigorous standardization of the plate preparation protocol, some variability between plates can occur. For each experiment, we correlated the number of migrated neutrophils from each well to the well’s initial TEER. Linear regression analysis showed that there was a very significant (p<0.0001) non-zero slope for the correlation between TEER and migrated neutrophil number in only the two plates with mean TEER 99.5 (+/- 41.4) and 206.3 (+/-48.9) and these two correlations also had the highest r^2^ values of 0.3988 and 0.2753, respectively. One plate had a significant but weak correlation with p = 0.001 and r^2^ = 0.1316. The remaining two plates had no significant correlation. These results suggest that TEER is related to migrated neutrophil number only when the mean plate TEER is at or below 206.3 Ω*cm^2^. In conjunction with this, we studied the effect of edge position on the number of migrated cells. The plates with mean TEER below 200 Ω*cm^2^ that demonstrated TEER-dependent edge effects also showed association between edge position and neutrophil number. Plate 3 did not exhibit TEER edge effects but did have neutrophil number edge effects. We speculate that this effect was not biological but rather was observed due to evaporation of edge wells during storage of the fixed cells before flow cytometry, concentrating the cells and creating a falsely higher value for edge wells during cytometry. Plate 5 experienced TEER and neutrophil edge effects, and we expect this was due to increased cell migration in wells with lower TEER due to edge-dependent differences in cell confluence, differentiation, or both. Control of plate evaporation during all stages of handling negates cell number edge effect in plates with no TEER-based edge effect.

---

**Supplemental Figure 1.**
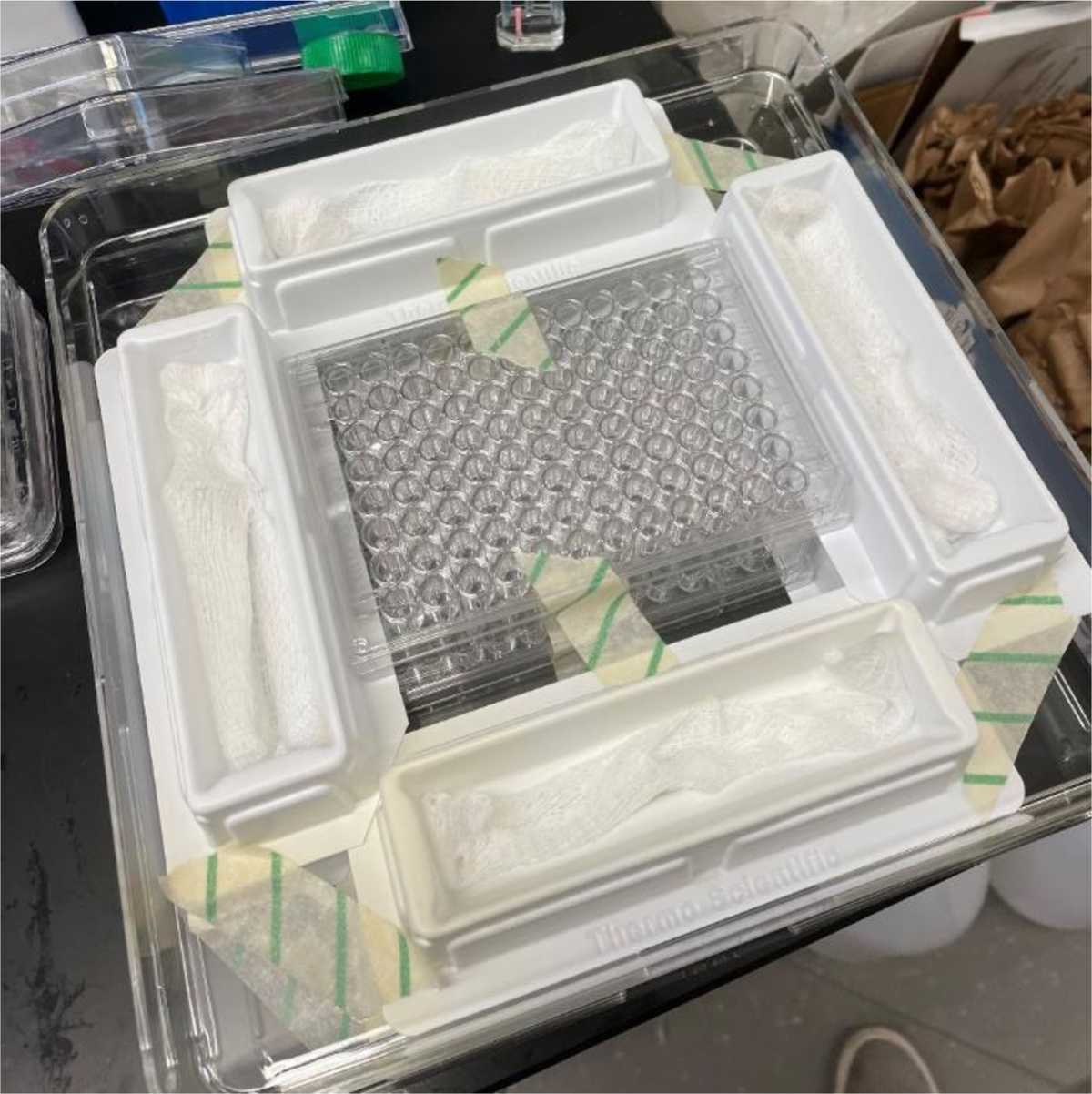
L-ABBA-96 humidity chamber to control edge effects.

**Supplemental Figure 2.**
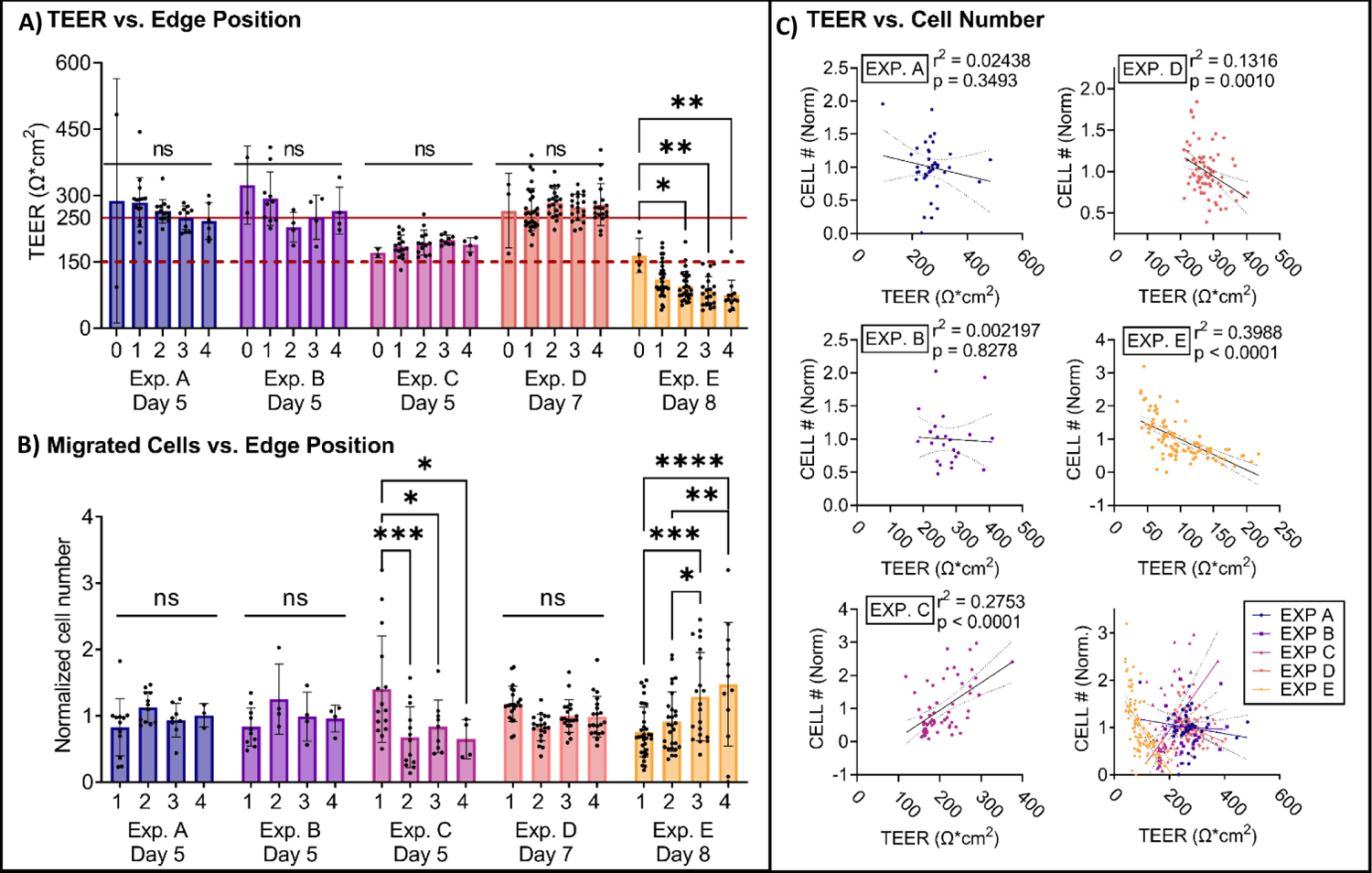
Migrated neutrophil number is free of TEER- and edge-dependence above a TEER threshold. In 5 independent Transwell plates cultured on different days, TEER of each well was measured immediately prior to neutrophil transmigration for 16 hours. Then, the number of migrated neutrophils was counted and correlated with initial TEER using simple linear regression followed by Wald’s test for slope significance (GraphPad Prism 9). **A)** TEER immediately prior to transmigration was independent of edge position unless the average plate TEER was below 150 Ω*cm² (red dashed line). **B)** From the same 5 independent Transwell plates, the number of migrated neutrophils was independent of edge position unless the average TEER was below 200 Ω*cm² (red solid line in **A)**). **C)** Migrated cell number was not strongly correlated with TEER (p > 0.05) when mean plate TEER exceeded approximately 200 Ω*cm². Strong correlations, defined as r² > 0.25 and p < 0.0001, had opposite slopes. Overall, no consistent trend between TEER and cell number was observed. n = 38, 24, 59, 79, and 96 wells for Experiments 1-5, respectively. All statistics were performed on GraphPad Prism 9 as follows. A, B: Two-way ANOVA with post-hoc Tukey’s t-test for multiple comparisons without matching. C: Simple linear regression with α = 0.05 and Wald test for significantly nonzero linear regression slope. ns = not significant, *p≤0.05, **p≤0.01, ***p≤0.001, ****p≤0.0001

**Supplemental Figure 3.**
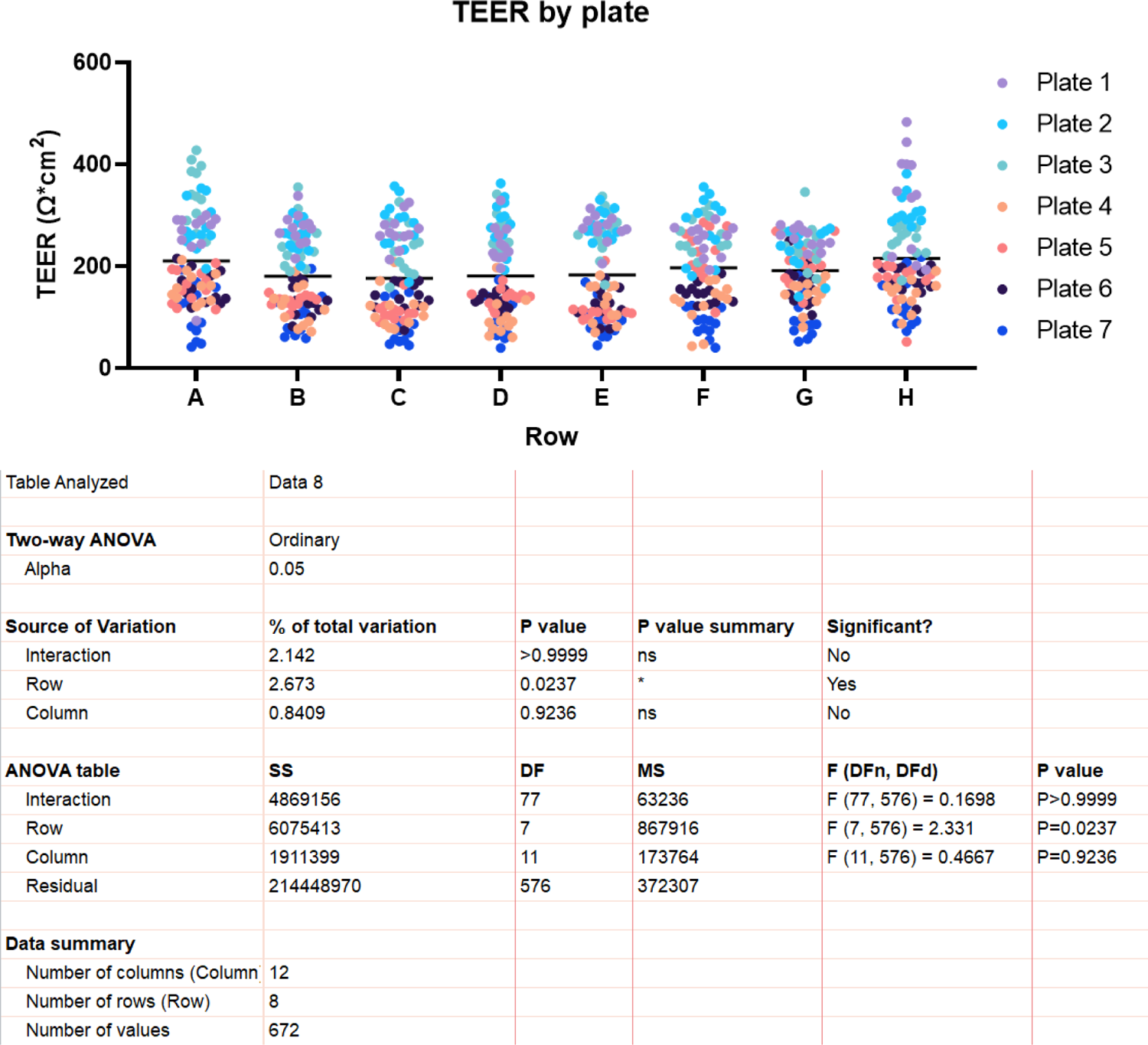
Plates with average TEER ∼200 Ω*cm^2^ show negligible row and column effects according to two-way ANOVA (Graphpad Prism 9).

**Supplemental Figure 4.**
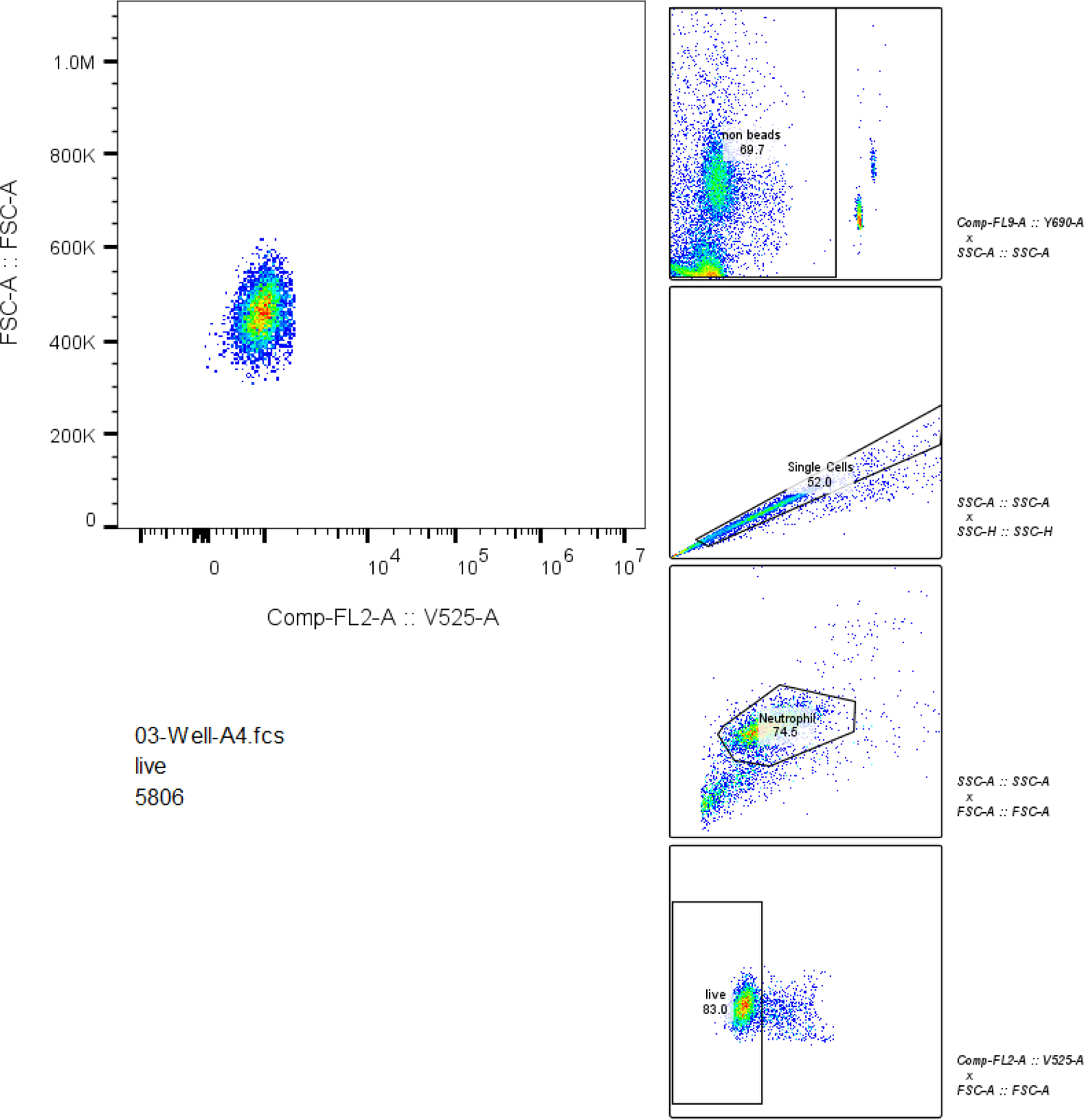
Gating strategy for flow cytometry.

**Supplemental Figure 5.**
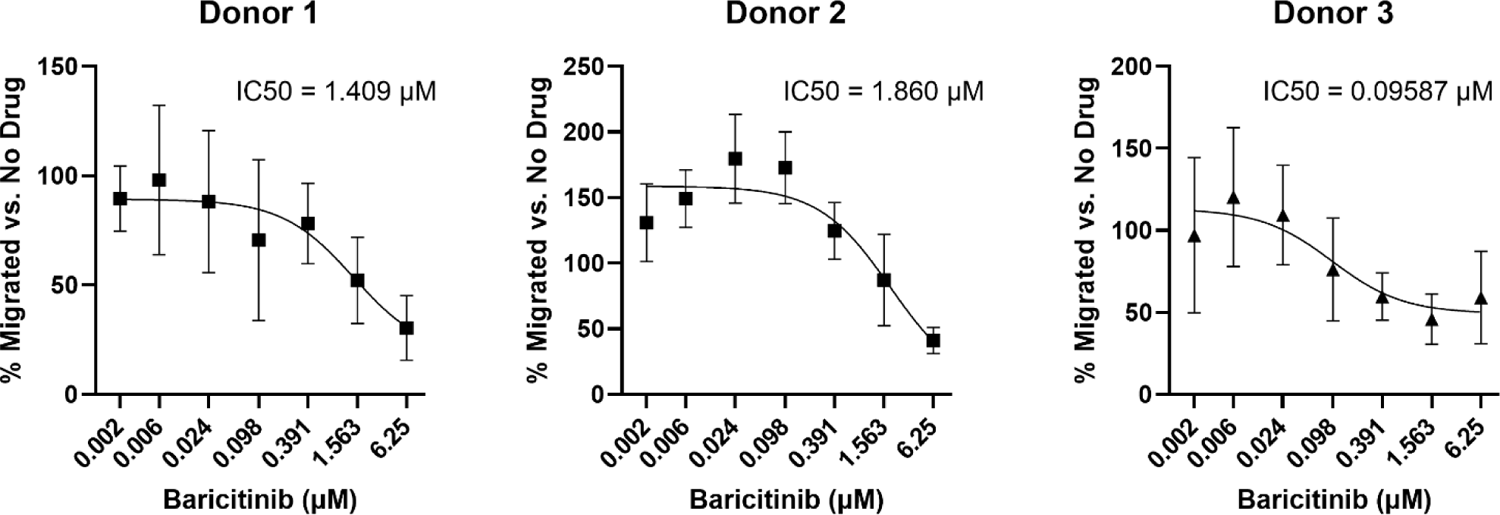
Donor dose-response neutrophil # with IC50s. Neutrophil number quantified by flow cytometry using counting beads. IC50 calculated using the standard 3-point log10 inhibitor dose-response logistic regression model in GraphPad Prism 9.

**Supplemental Figure 6.**
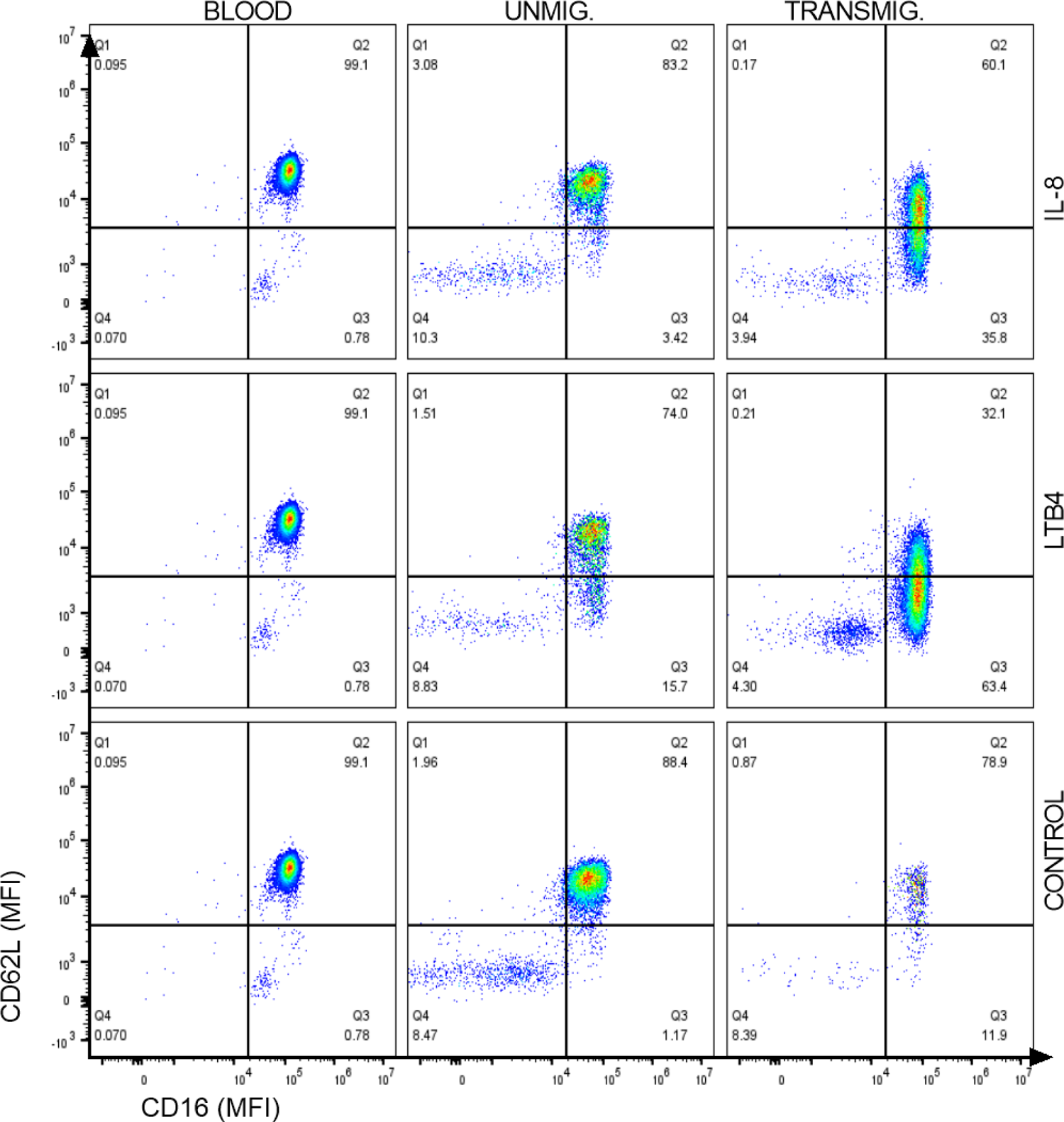
Representative plots showing the gating of CD62L and CD16 and shifts across various conditions.

**Supplemental Table 1.** Components of total variation in neutrophil response to IL-8 (Two-way ANOVA)

| Source of Variation | CD62L | CD66b | CD16 | CD63 | Mean |
| --- | --- | --- | --- | --- | --- |
| Condition x Donor | 3.237 | 12.74 | 12.79 | 14.25 | 10.75 ± 4.38 |
| Condition | 90.23 | 70.34 | 66.57 | 21.17 | 62.08 ± 25.27 |
| Donor | 6.392 | 12.72 | 19.22 | 56.08 | 23.6 ± 19.29 |

**Supplemental Table 2.** p values (Two-way ANOVA)

| Source of Variation | CD62L | CD66b | CD16 | CD63 |
| --- | --- | --- | --- | --- |
| Condition x Donor | **** | **** | **** | *** |
| Condition | **** | **** | **** | **** |
| Donor | **** | **** | **** | **** |

**Supplemental Table 3.** Flow panel for all experiments.

| Target | Color | Vendor | Cat # | Product Name | Antibody Type |
| --- | --- | --- | --- | --- | --- |
| CD63 | AF647 | BIOLEGEND | 353016 | Alexa Fluor® 647 anti-human CD63 | Mouse IgG1, κ |
| CD66B | PC7 | BIOLEGEND | 305116 | PE/Cyanine7 anti-human CD66b | Mouse IgM, κ |
| CD16 | APC-CY7 | BIOLEGEND | 302018 | APC/Cyanine7 anti-human CD16 | Mouse IgG1, κ |
| CD62L | BV421 | BIOLEGEND | 304828 | Brilliant Violet 421™ anti-human CD62L | Mouse IgG1, κ |
| Live/dead | KO525 | BIOLEGEND | 423101 | Zombie Aqua™ Fixable Viability Kit | N/A |

## References

1. Summers, C. Chasing the ‘Holy Grail’ – Modulating Neutrophils in Inflammatory Lung Disease. Am. J. Respir. Crit. Care Med. (2019) doi:10.1164/rccm.201902-0333ED.

2. Németh, T., Sperandio, M. & Mócsai, A. Neutrophils as emerging therapeutic targets. Nat. Rev. Drug Discov. 19, 253–275 (2020).

3. Cruz, M. A. et al. Nanomedicine platform for targeting activated neutrophils and neutrophil–platelet complexes using an α1-antitrypsin-derived peptide motif. Nat. Nanotechnol. 17, 1004–1014 (2022).

4. Amratia, D. A., Viola, H. & Ioachimescu, O. C. Glucocorticoid therapy in respiratory illness: bench to bedside. J. Investig. Med. (2022) doi:10.1136/jim-2021-002161.

5. Sinha, S. et al. Dexamethasone modulates immature neutrophils and interferon programming in severe COVID-19. Nat. Med. 28, 201–211 (2022).

6. Wang, Y. et al. Small-Molecule Modulators of Toll-like Receptors. Acc. Chem. Res. 53, 1046–1055 (2020).

7. Metzemaekers, M., Gouwy, M. & Proost, P. Neutrophil chemoattractant receptors in health and disease: double-edged swords. Cell. Mol. Immunol. 17, 433–450 (2020).

8. Othman, A., Sekheri, M. & Filep, J. G. Roles of neutrophil granule proteins in orchestrating inflammation and immunity. FEBS J. 289, 3932–3953 (2022).

9. Margraf, A., Lowell, C. A. & Zarbock, A. Neutrophils in acute inflammation: current concepts and translational implications. Blood 139, 2130–2144 (2022).

10. Stackowicz, J., Jönsson, F. & Reber, L. L. Mouse Models and Tools for the in vivo Study of Neutrophils. Front. Immunol. 10, (2020).

11. Patil, K. R. et al. Animal Models of Inflammation for Screening of Anti-inflammatory Drugs: Implications for the Discovery and Development of Phytopharmaceuticals. Int. J. Mol. Sci. 20, 4367 (2019).

12. Matsushima, K., Yang, D. & Oppenheim, J. J. Interleukin-8: An evolving chemokine. Cytokine 153, 155828 (2022).

13. Cesta, M. C. et al. The Role of Interleukin-8 in Lung Inflammation and Injury: Implications for the Management of COVID-19 and Hyperinflammatory Acute Respiratory Distress Syndrome. Front. Pharmacol. 12, (2022).

14. Irimia, D. & Wang, X. Inflammation-on-a-Chip: Probing the Immune System Ex Vivo. Trends Biotechnol. 36, 923–937 (2018).

15. Han, S. et al. A versatile assay for monitoring in vivo -like transendothelial migration of neutrophils. Lab. Chip 12, 3861–3865 (2012).

16. Huh, D. et al. Reconstituting Organ-Level Lung Functions on a Chip. Science 328, 1662–1668 (2010).

17. Chew, K. et al. A protocol for high-throughput screening for immunomodulatory compounds using human primary cells. STAR Protoc. 4, 102405 (2023).

18. Lv, D. et al. A novel cell-based assay for dynamically detecting neutrophil extracellular traps-induced lung epithelial injuries. Exp. Cell Res. 394, 112101 (2020).

19. Vogt, K. L., Summers, C., Chilvers, E. R. & Condliffe, A. M. Priming and de-priming of neutrophil responses in vitro and in vivo. Eur. J. Clin. Invest. 48, e12967 (2018).

20. Kumar, S. D., Krishnamurthy, K., Manikandan, J., Pakeerappa, P. N. & Pushparaj, P. N. Deciphering the key molecular and cellular events in neutrophil transmigration during acute inflammation. Bioinformation 6, 111–114 (2011).

21. Hinman, S. S. et al. Microphysiological system design: simplicity is elegance. Curr. Opin. Biomed. Eng. 13, 94–102 (2020).

22. Probst, C., Schneider, S. & Loskill, P. High-throughput organ-on-a-chip systems: Current status and remaining challenges. Curr. Opin. Biomed. Eng. 6, 33–41 (2018).

23. Kamm, R. D. et al. Perspective: The promise of multi-cellular engineered living systems. APL Bioeng. 2, 040901 (2018).

24. Viola, H. et al. Microphysiological systems modeling acute respiratory distress syndrome that capture mechanical force-induced injury-inflammation-repair. APL Bioeng. 3, 041503 (2019).

25. Huh, D. et al. Acoustically detectable cellular-level lung injury induced by fluid mechanical stresses in microfluidic airway systems. Proc. Natl. Acad. Sci. 104, 18886–18891 (2007).

26. Viola, H. L. et al. Liquid plug propagation in computer-controlled microfluidic airway-on-a-chip with semi-circular microchannels. Lab. Chip (2024) doi:10.1039/D3LC00957B.

27. Douville, N. J. et al. Combination of fluid and solid mechanical stresses contribute to cell death and detachment in a microfluidic alveolar model. Lab Chip 11, 609–619 (2011).

28. C. Mejías, J. R. Nelson, M., Liseth, O. & Roy, K. A 96-well format microvascularized human lung-on-a-chip platform for microphysiological modeling of fibrotic diseases. Lab. Chip 20, 3601–3611 (2020).

29. Plebani, R. et al. Modeling pulmonary cystic fibrosis in a human lung airway-on-a-chip. J. Cyst. Fibros. (2021) doi:10.1016/j.jcf.2021.10.004.

30. Nawroth, J. C. et al. A Microengineered Airway Lung Chip Models Key Features of Viral-induced Exacerbation of Asthma. Am. J. Respir. Cell Mol. Biol. 63, 591–600 (2020).

31. Nazari, H. et al. Advances in TEER measurements of biological barriers in microphysiological systems. Biosens. Bioelectron. 234, 115355 (2023).

32. Ekpenyong, A. E., Toepfner, N., Chilvers, E. R. & Guck, J. Mechanotransduction in neutrophil activation and deactivation. Biochim. Biophys. Acta BBA - Mol. Cell Res. 1853, 3105–3116 (2015).

33. Shive, M. S., Salloum, M. L. & Anderson, J. M. Shear stress-induced apoptosis of adherent neutrophils: A mechanism for persistence of cardiovascular device infections. Proc. Natl. Acad. Sci. 97, 6710–6715 (2000).

34. Candarlioglu, P. L. et al. Organ-on-a-chip: current gaps and future directions. Biochem. Soc. Trans. 50, 665–673 (2022).

35. Benam, K. H. et al. Small airway-on-a-chip enables analysis of human lung inflammation and drug responses in vitro. Nat. Methods 13, 151–157 (2016).

36. Si, L. et al. A human-airway-on-a-chip for the rapid identification of candidate antiviral therapeutics and prophylactics. *Nat*. Biomed. Eng. 5, 815–829 (2021).

37. Forrest, O. A. et al. Frontline Science: Pathological conditioning of human neutrophils recruited to the airway milieu in cystic fibrosis. J. Leukoc. Biol. 104, 665– 675 (2018).

38. Forrest, O. A. et al. Neutrophil-derived extracellular vesicles promote feed-forward inflammasome signaling in cystic fibrosis airways. J. Leukoc. Biol. 112, 707–716 (2022).

39. Maas, S. L. & van der Vorst, E. P. C. In Vitro (Trans)Migration Experiment Using Chemokines as Stimulatory Factor. in Chemokine-Glycosaminoglycan Interactions: Methods and Protocols (ed. Lucas, A. R.) 77–87 (Springer US, New York, NY, 2023). doi:10.1007/978-1-0716-2835-5_7.

40. Dobosh, B., Giacalone, V. D., Margaroli, C. & Tirouvanziam, R. Mass production of human airway-like neutrophils via transmigration in an organotypic model of human airways. STAR Protoc. 2, 100892 (2021).

41. Viola, H. et al. A High-Throughput Distal Lung Air–Blood Barrier Model Enabled By Density-Driven Underside Epithelium Seeding. Adv. Healthc. Mater. 2100879 (2021) doi:10.1002/adhm.202100879.

42. Gschwandtner, M. et al. Glycosaminoglycans are important mediators of neutrophilic inflammation *in vivo*. Cytokine 91, 65–73 (2017).

43. Alflen, A. et al. ADAMTS-13 regulates neutrophil recruitment in a mouse model of invasive pulmonary aspergillosis. Sci. Rep. 7, 7184 (2017).

44. Adams, W., Espicha, T. & Estipona, J. Getting Your Neutrophil: Neutrophil Transepithelial Migration in the Lung. Infect. Immun. 89, e00659–20.

45. Lin, W.-C. & Fessler, M. B. Regulatory mechanisms of neutrophil migration from the circulation to the airspace. Cell. Mol. Life Sci. 78, 4095–4124 (2021).

46. Rubin, R. Baricitinib Is First Approved COVID-19 Immunomodulatory Treatment. JAMA 327, 2281 (2022).

47. Bronte, V. et al. Baricitinib restrains the immune dysregulation in patients with severe COVID-19. J. Clin. Invest. 130, 6409–6416 (2020).

48. Condliffe, A. M., Chilvers, E. R., Haslett, C. & Dransfield, I. Priming differentially regulates neutrophil adhesion molecule expression/function. Immunology 89, 105– 111 (1996).

49. Miralda, I., Uriarte, S. M. & McLeish, K. R. Multiple Phenotypic Changes Define Neutrophil Priming. Front. Cell. Infect. Microbiol. 7, (2017).

50. Mazzitelli, I. et al. Immunoglobulin G Immune Complexes May Contribute to Neutrophil Activation in the Course of Severe Coronavirus Disease 2019. J. Infect. Dis. 224, 575–585 (2021).

51. Monboisse, J.-C., Garnotel, R., Randoux, A., Dufer, J. & Borel, J.-P. Adhesion of Human Neutrophils to and Activation by Type-I Collagen Involving a β2 Integrin. J. Leukoc. Biol. 50, 373–380 (1991).

52. Giacalone, V. D., Margaroli, C., Mall, M. A. & Tirouvanziam, R. Neutrophil Adaptations upon Recruitment to the Lung: New Concepts and Implications for Homeostasis and Disease. Int. J. Mol. Sci. 21, 851 (2020).

53. Mansoury, M., Hamed, M., Karmustaji, R., Al Hannan, F. & Safrany, S. T. The edge effect: A global problem. The trouble with culturing cells in 96-well plates. Biochem. Biophys. Rep. 26, 100987 (2021).

54. Wilson, G. A. et al. Human-specific epigenetic variation in the immunological Leukotriene B4 Receptor (LTB4R/BLT1) implicated in common inflammatory diseases. Genome Med. 6, 19 (2014).

55. Ivetic, A. Signals regulating L-selectin-dependent leucocyte adhesion and transmigration. Int. J. Biochem. Cell Biol. 45, 550–555 (2013).

56. Ivetic, A., Hoskins Green, H. L. & Hart, S. J. L-selectin: A Major Regulator of Leukocyte Adhesion, Migration and Signaling. Front. Immunol. 10, (2019).

57. Middelhoven, P. J., Ager, A., Roos, D. & Verhoeven, A. J. Involvement of a metalloprotease in the shedding of human neutrophil FcγRIIIB. FEBS Lett. 414, 14– 18 (1997).

58. Middelhoven, P. J., van Buul, J. D., Kleijer, M., Roos, D. & Hordijk, P. L. Actin Polymerization Induces Shedding of FcγRIIIb (CD16) from Human Neutrophils. Biochem. Biophys. Res. Commun. 255, 568–574 (1999).

59. Tosi, M. F. & Zakem, H. Surface expression of Fc gamma receptor III (CD16) on chemoattractant-stimulated neutrophils is determined by both surface shedding and translocation from intracellular storage compartments. https://www.jci.org/articles/view/115882/pdf<x> (1992) doi:10.1172/JCI115882.

60. Moulding, D. A., Hart, C. A. & Edwards, S. W. Regulation of neutrophil FcγRIIIb (CD16) surface expression following delayed apoptosis in response to GM-CSF and sodium butyrate. J. Leukoc. Biol. 65, 875–882 (1999).

61. Fortunati, E., Kazemier, K. M., Grutters, J. C., Koenderman, L. & Van den Bosch, V. J. M. M. Human neutrophils switch to an activated phenotype after homing to the lung irrespective of inflammatory disease. Clin. Exp. Immunol. 155, 559–566 (2009).

62. Ley, K. Integration of inflammatory signals by rolling neutrophils. Immunol. Rev. 186, 8–18 (2002).

63. Jorgensen, S. C. J., Tse, C. L. Y., Burry, L. & Dresser, L. D. Baricitinib: A Review of Pharmacology, Safety, and Emerging Clinical Experience in COVID-19. Pharmacother. J. Hum. Pharmacol. Drug Ther. 40, 843–856 (2020).

64. McGonagle, D., Sharif, K., O’Regan, A. & Bridgewood, C. The Role of Cytokines including Interleukin-6 in COVID-19 induced Pneumonia and Macrophage Activation Syndrome-Like Disease. Autoimmun. Rev. 19, 102537 (2020).

65. Meduri, G. U. et al. Inflammatory Cytokines in the BAL of Patients With ARDS: Persistent Elevation Over Time Predicts Poor Outcome. Chest 108, 1303–1314 (1995).

66. Mukaida, N. Pathophysiological roles of interleukin-8/CXCL8 in pulmonary diseases. Am. J. Physiol.-Lung Cell. Mol. Physiol. 284, L566–L577 (2003).

67. Kubo, S. et al. Janus Kinase Inhibitor Baricitinib Modulates Human Innate and Adaptive Immune System. Front. Immunol. 9, 1510 (2018).

68. Ronit, A. et al. Compartmental immunophenotyping in COVID-19 ARDS: A case series. J. Allergy Clin. Immunol. 147, 81–91 (2021).

69. Huang, W. et al. The Inflammatory Factors Associated with Disease Severity to Predict COVID-19 Progression. J. Immunol. 206, 1597–1608 (2021).

70. Hoang, T. N. et al. Baricitinib treatment resolves lower-airway macrophage inflammation and neutrophil recruitment in SARS-CoV-2-infected rhesus macaques. Cell 184, 460–475.e21 (2021).

71. Barilli, A. et al. The JAK1/2 Inhibitor Baricitinib Mitigates the Spike-Induced Inflammatory Response of Immune and Endothelial Cells In Vitro. Biomedicines 10, 2324 (2022).

72. Henkels, K. M., Frondorf, K., Gonzalez-Mejia, M. E., Doseff, A. L. & Gomez-Cambronero, J. IL-8-induced neutrophil chemotaxis is mediated by Janus kinase 3 (JAK3). FEBS Lett. 585, 159–166 (2011).

73. Wang, Z. & Chan, E. C. Y. Physiologically-Based Pharmacokinetic Modelling to Investigate Baricitinib and Tofacitinib Dosing Recommendations for COVID-19 in Geriatrics. Clin. Pharmacol. Ther. 112, 291–296 (2022).

74. Pillay, J. et al. A subset of neutrophils in human systemic inflammation inhibits T cell responses through Mac-1. J. Clin. Invest. 122, 327–336 (2012).

75. Tak, T. et al. Human CD62Ldim neutrophils identified as a separate subset by proteome profiling and in vivo pulse-chase labeling. Blood 129, 3476–3485 (2017).

76. Spijkerman, R. et al. An increase in CD62Ldim neutrophils precedes the development of pulmonary embolisms in COVID-19 patients. Scand. J. Immunol. 93, e13023 (2021).

77. de Formiga, R. O., et al. Cytosolic PCNA interacts with S100A8 and controls an inflammatory subset of neutrophils in COVID-19. 2022.10.12.22280984 Preprint at 10.1101/2022.10.12.22280984 (2022).

78. De Rose, V. et al. Circulating Adhesion Molecules in Cystic Fibrosis. Am. J. Respir. Crit. Care Med. 157, 1234–1239 (1998).

79. Declercq, M., Treps, L., Carmeliet, P. & Witters, P. The role of endothelial cells in cystic fibrosis. J. Cyst. Fibros. 18, 752–761 (2019).

80. Tirouvanziam, R., Khazaal, I. & Péault, B. Primary inflammation in human cystic fibrosis small airways. Am. J. Physiol.-Lung Cell. Mol. Physiol. 283, L445–L451 (2002).

81. Martin, C. et al. Specific circulating neutrophils subsets are present in clinically stable adults with cystic fibrosis and are further modulated by pulmonary exacerbations. Front. Immunol. 13, (2022).

82. Kamp, V. M. et al. Human suppressive neutrophils CD16bright/CD62Ldim exhibit decreased adhesion. J. Leukoc. Biol. 92, 1011–1020 (2012).

83. Oshika, E. et al. Glucocorticoid-Induced Effects on Pattern Formation and Epithelial Cell Differentiation in Early Embryonic Rat Lungs. Pediatr. Res. 43, 305–314 (1998).

84. Lin, Z. et al. Clinical efficacy and adverse events of baricitinib treatment for coronavirus disease-2019 (COVID-19): A systematic review and meta-analysis. J. Med. Virol. 94, 1523–1534 (2022).

85. Kaiser, R. et al. Self-sustaining IL-8 loops drive a prothrombotic neutrophil phenotype in severe COVID-19. JCI Insight 6, e150862.

86. Williams, M. A. & Solomkin, J. S. Integrin-mediated signaling in human neutrophil functioning. J. Leukoc. Biol. 65, 725–736 (1999).

87. Hazeldine, J. et al. Prehospital immune responses and development of multiple organ dysfunction syndrome following traumatic injury: A prospective cohort study. PLOS Med. 14, e1002338 (2017).

88. Hao, S., Andersen, M. & Yu, H. Detection of immune suppressive neutrophils in peripheral blood samples of cancer patients. Am. J. Blood Res. 3, 239–245 (2013).

89. Hassani, M. et al. On the origin of low-density neutrophils. J. Leukoc. Biol. 107, 809– 818 (2020).

## REFERENCES

1. Kidney, J. C. & Proud, D. Neutrophil Transmigration across Human Airway Epithelial Monolayers. Am. J. Respir. Cell Mol. Biol. 23, 389–395 (2000).

